# Activity-dependent capture reveals brain-wide signatures of isoflurane anesthesia-induced unconsciousness

**DOI:** 10.64898/2025.12.02.691631

**Authors:** Minke H.C. Nota, Lukas K. MacMillen, Hallie Lazaro, Jaidyn O. Gaur, Ian Campuzano, Carlie Neiswanger, Sam A. Golden, Mitra Heshmati

## Abstract

**Background:** General anesthesia is commonly used to produce unconsciousness across species, though the underlying neural substrates remain poorly understood. Here we test the hypothesis that isoflurane anesthesia produces unconsciousness by targeting discrete cell types and neural circuits distributed brain-wide, rather than by brain-wide non-specific binding or binding exclusively within single brain regions.

**Methods:** We take advantage of a transgenic mouse system that expresses chemogenetic designer receptors exclusively activated by a designer drug (DREADDs) in an inducible Cre-dependent manner driven by Fos immediate early gene expression. Since Fos peaks with metabolic activity, we use this system to insert DREADDs brain-wide into neurons that are active under isoflurane anesthesia. We then test the effects of chemogenetic manipulation of brain-wide anesthesia-activated neural ensembles on behavior, in the absence of isoflurane.

**Results:** Using iDisco+ intact brain clearing and light sheet microscopy, we describe brain-wide expression of the captured neurons revealing sparse, heterogeneous and spatially distributed cells. We quantify dense labeling across mesolimbic pathways that include the amygdala, hypothalamus, thalamus, and hindbrain nuclei, implicating these regions as candidate mediators of the anesthetic state. Chemogenetic manipulation of the brain-wide activated neural ensembles reproduces key components of anesthesia, including immobility and thermal anti-nociception. We observe significantly increased isoflurane sensitivity, though not a complete loss of consciousness as measured by the righting reflex, suggesting alternative mechanisms contribute to unconsciousness that are not captured by specific neural signatures.

**Conclusions:** We provide causal evidence that isoflurane anesthesia engages discrete, distributed brain-wide circuitry, recapitulating dissociable components of general anesthesia.

**Significance Statement:** Over 300 million people worldwide undergo general anesthesia each year, yet the neural circuit mechanisms that produce unconsciousness remain unresolved. This fundamental gap limits the development of safer anesthetic strategies and hinders efforts to understand long-term cognitive consequences. Here we define the intact brain-wide neural signatures of isoflurane anesthesia-induced unconsciousness using a transgenic mouse system. We capture brain-wide neurons that are activated by isoflurane, manipulate those ensembles using chemogenetic tools and characterize their spatial representations at the cellular resolution using light sheet microscopy. Together, our data provides causal evidence that anesthesia engages discrete, distributed brain-wide circuitry, offering a systems-level framework for understanding how unconsciousness is generated.

## Introduction

General anesthesia, which produces a reversible loss of consciousness, is a useful tool to study mechanisms of consciousness, particularly due to its evolutionarily conserved efficacy across phylogenetic kingdoms (1). Inhaled anesthetics such as isoflurane alter brain-wide functional connectivity in mice (2), non-human primates (3), and humans (4), and require the simultaneous recruitment of multiple neural networks for anesthetic effects (3, 4). General anesthesia is composed of distinct behavioral outcomes, such as loss of consciousness, immobility, and loss of pain perception or analgesia (5). Studies of discrete neural circuits have been effective at dissecting individual behavioral outcomes. These include the stimulation of thalamocortical neuronal populations (6, 7) or subcortical arousal nuclei (8, 9) to promote wakefulness, as well as modulation of thalamocortical connectivity to induce sedation and anesthetic-like EEG signatures (10). Stimulation of the preoptic nucleus induces sedation (11, 12) whereas lesions cause resistance to anesthesia-induced unconsciousness (13). Studies have further shown that anterior cingulate cortex stimulation inhibits spinal cord responses to noxious stimuli (14) and activation of the lateral parabrachial nucleus alleviates neuropathic pain (15). While these studies localize discrete circuit nodes with the ability to produce aspects of general anesthesia, they do not recapitulate the full scope of anesthetic outcomes. We hypothesize that the recapitulation of all anesthetic outcomes by neural circuits requires manipulating the global network.

Fluorescent light sheet microscopy combined with whole brain immunolabeling and clearing methods now allows for the capture of discrete activated neural circuitry across the intact brain. Genetic whole brain neural activity capture using Targeted Recombination in Active Populations (TRAP) is ideally suited to achieve detailed maps of anesthesia-induced unconsciousness within the intact system. With this method, behaviorally activated neuronal ensembles are co-opted for artificial stimulation or inhibition via genetic tools (17). Here we aim to synthetically capture brain-wide isoflurane-activated neural ensembles using a transgenic mouse system to better define neural circuit signatures of anesthesia-induced unconsciousness within the intact brain-wide network.

We crossed heterozygous R26-LSL-Gq-DREADD^+/-^ mice (18, 19) with homozygous Fos^2A-iCreER^ (Fos-TRAP2) mice (17), generating R26-LSL-Gq-DREADD^+^xFos^2A-iCreER+^ (Gq+) and R26-LSL-null-xFos^2A-iCreER+^ (*null*) mice (**Fig 1A**). As a result, all offspring have a FosTRAP2 genotype, but only Gq+ mice express the excitatory Gq-DREADD protein in an activity-dependent, tamoxifen-inducible manner. This allows for chemogenetic neural activity capture and synthetic stimulation in the awake state in the Gq+ mice. We hypothesize that this transgenic mouse facilitates improved and unbiased capture of brain-wide anesthesia-activated neurons, providing an alternative preclinical system for studying circuit and cell-type-specific unconsciousness circuitry across the intact brain. As a complement to chemogenetic functional manipulations, we further hypothesize that DREADD labeling within this transgenic system enables mapping the brain-wide cellular architecture of anesthesia circuitry. Together, these experiments test the overarching hypothesis that specific brain-wide neural circuit substrates exist to generate anesthesia-induced behavioral sedation, which we operationally define here as loss of the righting reflex, with further assays quantifying locomotion, motor coordination, and nociception that are additional behavioral outcomes affected by anesthesia.

**Fig 1.**
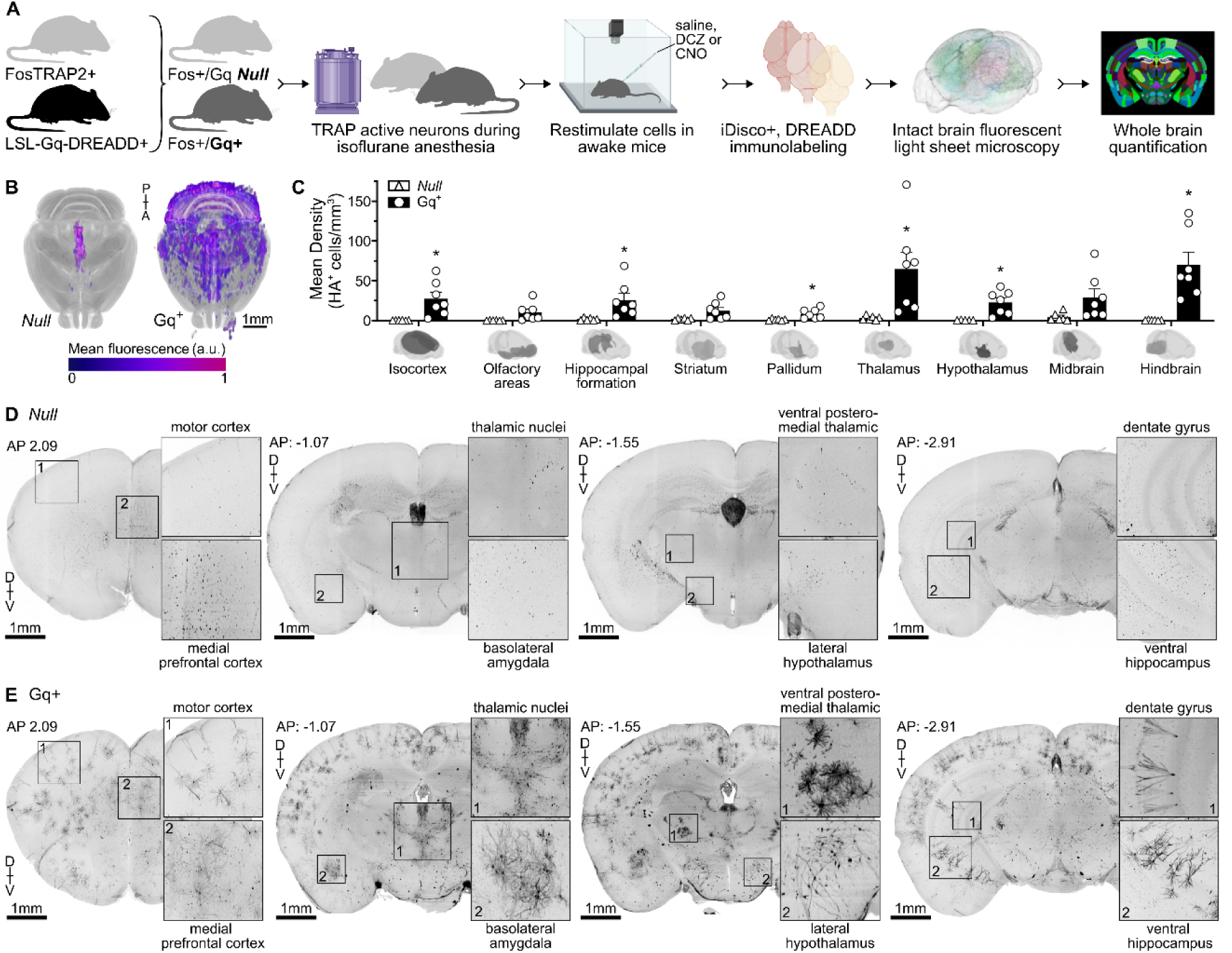
Activity-dependent neural capture across the intact brain. **A)** Breeding strategy and experiment timeline: Heterozygous R26-LSL-Gq-DREADD mice were crossed with homozygous Fos2A-iCreER (FosTRAP2) mice. Adult male and female offspring (heterozygous FosTRAP2-Gq^+^ and *null* FosTRAP2 littermates) underwent 3-hour isoflurane anesthesia (1.2-1.3%) exposure, with intraperitoneal 4-hydroxytamoxifen (4-OHT) given halfway through to induce activity-dependent DREADD labeling (TRAP). The behavioral effects of subsequent DREADD stimulation with clozapine-N-oxide (CNO) or saline were assessed using righting reflex, rotarod, modified open field test, and warm water tail withdrawal. Intact brains were cleared and immunolabeled for HA-tagged DREADD using a modified iDisco+ pipeline. Brains were imaged using a light sheet microscope and HA expression was quantified with ClearMap. **B)** Whole brain renders of HA expression in *null* and Gq^+^ mice. **C)** Quantification of HA^+^ DREADD-expressing cells across major brain divisions. **D)** *Null* mice (n=6) do not show neuronal labeling across the brain. **E)** In Gq^+^ mice (n=7), neural expression with detailed morphologic labeling is visualized across spatially distributed brain regions, including the motor and somatosensory cortex, medial prefrontal cortex, nucleus accumbens, thalamic nuclei, lateral hypothalamus, basolateral amygdala, and hippocampal subregions (dentate gyrus, CA1, subiculum). See also **Movie S1** and **Fig S1**. For detailed statistics see **Table S1**.

## Results

### Brain-wide analysis after chemogenetic capture identifies spatially distributed isoflurane-activated neurons

After registering intact brains to the Unified mouse brain atlas (20) and quantifying subregion DREADD expression with ClearMap (22), we found that Gq+ mice show significantly higher overall DREADD expression than *null* littermates, which do not show labeling across the brain (**Fig 1B–C**, **Movie S1**, **Fig S1**). Of 917 minor subregions analyzed, 272 (∼30%) are significantly enriched in Gq+ mice, and every one of the nine major anatomical divisions contained enriched subregions (**Fig 1C**, **Table S1**). Isoflurane-activated ensembles are therefore neither confined to a single locus nor uniformly distributed. DREADD expression forms a pattern that both recapitulates established anesthesia-relevant nuclei and extends into regions not typically emphasized in circuit-level studies of anesthesia. Below we highlight the most prominent regions across both categories (full subregion-level statistics are provided in **Table S1)**.

#### Recapitulation of established anesthesia- and arousal-relevant circuitry

DREADD expression is prominent across regions with well-documented roles in regulating consciousness, arousal, and anesthetic sensitivity. Within the hypothalamus, we observe dense labeling in the ventrolateral preoptic nucleus, a sleep-promoting structure directly activated by volatile anesthetics (13, 21, 22). We also observe expression within the lateral hypothalamic zone, anterior hypothalamic area, and ventral zona incerta. The thalamus has significant expression spanning the intralaminar, anterior, lateral, and posterior nuclear groups, the mediodorsal nucleus, and both medial and lateral geniculate nuclei. This pattern is consistent with thalamic regulation of cortical arousal and the disruption of thalamocortical connectivity reported during anesthesia (23–26). In the hindbrain, labeling is densest in structures implicated in arousal and antinociception, including the superior and lateral parabrachial nuclei, periaqueductal gray, median raphe, reticular formation, and pontine nuclei. The midbrain shows expression in subregions including the ventral tegmental area, substantia nigra (reticular part), and interpeduncular nucleus. Together, these regions form a recognizable subcortical scaffold for anesthesia-induced state transitions.

#### Engagement of thalamocortical and limbic circuits linked to behavioral outcomes of anesthesia

A second cluster of DREADD expression spans cortical and limbic structures that map onto the behavioral effects we observed downstream of chemogenetic stimulation. Across the isocortex, labeling is distributed in multiple layers of primary and secondary motor and somatosensory cortices, cingulate areas, and orbital, insular, parietal, and temporal association regions. Striatal enrichment is concentrated in the nucleus accumbens, a structure implicated in modulation of conscious state and pain (27–33), with additional labeling in the olfactory tubercle, caudoputamen, and amygdalostriatal transition zones. Limbic enrichment extends to the basolateral and cortical amygdaloid nuclei and the anterior amygdaloid area, regions central to descending antinociception (34–36).

#### Less established regions

Several patterns of DREADD expression are unexpected based on the existing anesthesia literature. Within the hippocampal formation, labeling is concentrated in subicular and parahippocampal structures (parasubiculum, dorsal and ventral subiculum, subiculum transition area, and caudomedial and dorsolateral entorhinal cortex), rather than in CA1 or CA3 pyramidal subfields, with the notable exception of the CA2 pyramidal layer. Primary and secondary sensory cortices outside the somatosensory system, including auditory and visual cortical layers, are also enriched. Finally, the cerebellum shows dense labeling in granular and molecular layers of the cerebellar cortex and the lateral medial cerebellar nucleus. Olfactory areas (anterior olfactory nucleus, piriform layer 3, nucleus of the lateral olfactory tract, endopiriform nucleus, dorsal peduncular cortex) and brainstem sensorimotor nuclei (motor and principal sensory trigeminal, facial, vestibular, cochlear, solitary, and lateral lemniscus nuclei, periolivary regions, medullary structures) are additional areas with significant DREADD expression.

### Chemogenetic stimulation increases sensitivity to loss of consciousness

Loss of consciousness is one of the defining characteristics of general anesthesia (4, 5). In mice, loss of righting reflex (LORR) is an established behavioral proxy for loss of consciousness (37–40). After chemogenetic capture, we assessed the contribution of brain-wide anesthesia-activated neuronal ensembles to loss of consciousness when chemogenetically stimulated in the awake state. We used deschloroclozapine (DCZ) as our primary DREADD agonist, as it is highly specific for the synthetic Gq-DREADD receptor and has a half-life of approximately 30-min, allowing for tight temporal coupling to behavioral assessment (41). In alternating trials, mice received 0.1mg/kg DCZ, or saline vehicle control, and there was not immediately an observable LORR. We then assessed sensitivity to isoflurane after chemogenetic stimulation. Mice received isoflurane in an incrementally increasing stepwise concentration over 10 min until LORR was observed, and the latency to return of righting reflex (RORR) after isoflurane exposure was recorded (**Fig 2A**). *Null* littermate controls did not show a change in LORR or RORR (**Fig 2B**). Gq^+^ mice displayed LORR at a significantly lower isoflurane concentration after DCZ injection, compared to saline (**Fig 2C**). Following isoflurane exposure, Gq^+^ mice showed a significantly increased latency to RORR (**Fig 2C**). See **Table S2** for detailed statistics.

**Fig 2.**
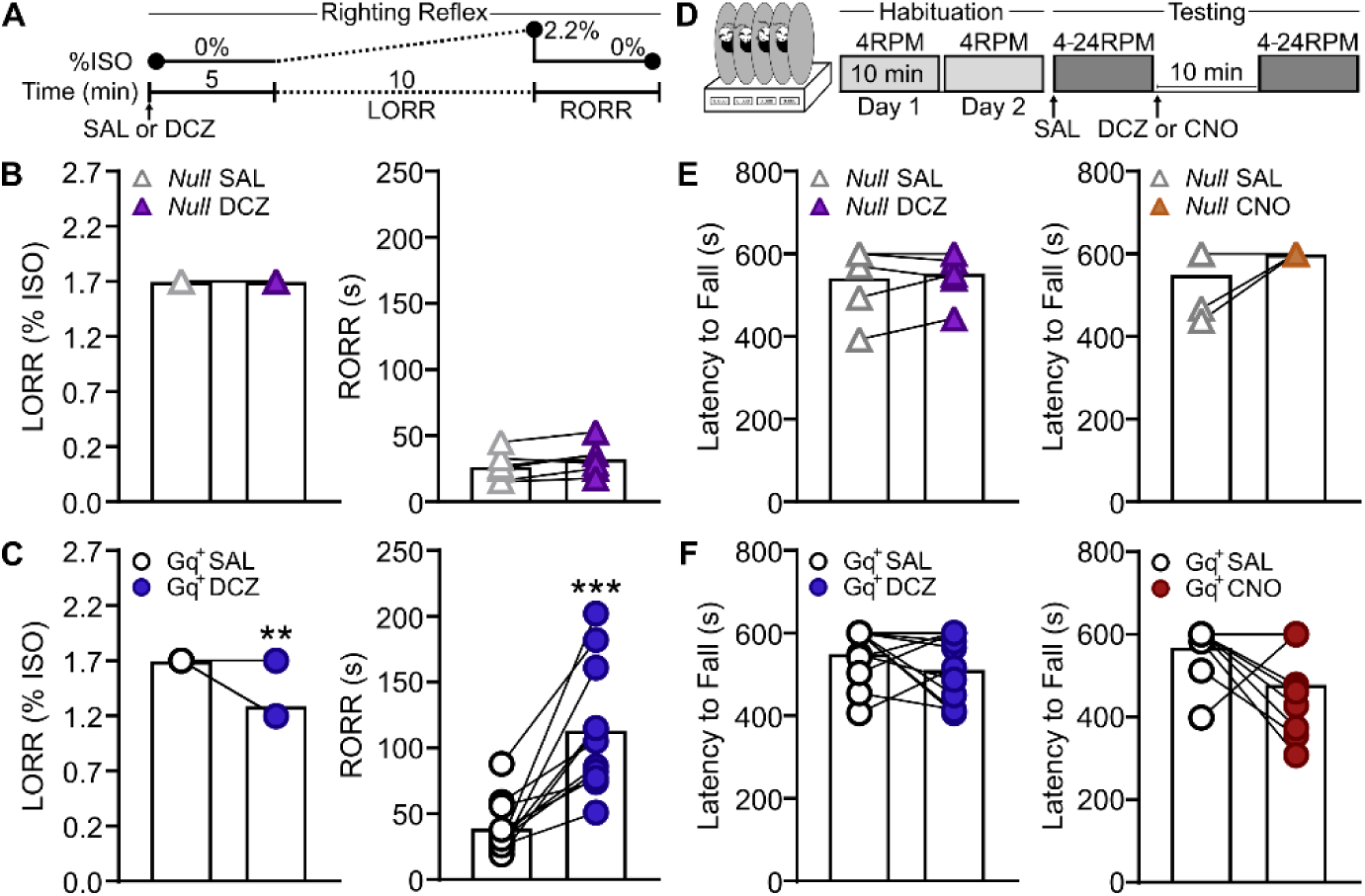
Chemogenetic stimulation increases sensitivity to isoflurane without significant effects on coordinated movement. **A)** Righting reflex testing: Mice underwent stepwise isoflurane induction from 0.0-2.2% (initial increment 0.2%, subsequent increments 0.5%). Each step was maintained for 2 min and the isoflurane concentration at which loss of righting reflex (LORR) occurred was recorded. After 2 min at 2.2%, isoflurane was turned off. Mice were placed in a supine position and the latency to return of righting reflex (RORR) was measured. **B)** In *null* mice (n=6), DCZ did not increase sensitivity to isoflurane or the latency to RORR. **C)** In Gq^+^ mice (n=11), DCZ increased the sensitivity to isoflurane and increased the latency to RORR to a much greater degree compared to *null* littermates. **D)** Rotarod habituation and testing: Mice were habituated to a rotarod twice for 10 min at 4RPM and subsequently tested at gradually increasing speed (4-24RPM over 10 min) either directly following saline injection or 10min following DCZ (0.1mg/kg) or CNO injection (5mg/kg). **E-F)** DCZ and CNO did not significantly affect coordination in *null* (n=6) or Gq^+^ (n=11) mice. Error bars represent standard error of the mean. **p<0.01, ***p<0.001 vs. saline. For detailed statistics see **Table S2**.

### Chemogenetic stimulation does not alter coordinated movement on the rotarod

To confirm that the effects on righting reflex were not caused by generalized motor disruption, we next assessed the effect of chemogenetic stimulation of isoflurane-activated neural circuitry on coordinated locomotion in the rotarod test. Gq^+^ mice and *null* littermate controls were habituated to a rotarod and subsequently tested at gradually increasing speed following saline or DCZ (0.1mg/kg) injections (**Fig 2D**). In both *null* and Gq^+^ mice, there were no significant effects of DCZ on the latency to fall compared to saline (**Fig 2E-F**). To confirm that the absence of effects was not due to the fast pharmacokinetics of DCZ, we also repeated testing with clozapine-n-oxide (CNO, 5mg/kg). CNO remains the most widely used DREADD agonist in the field, though studies show CNO can be reverse-metabolized to clozapine which may produce off-target behavioral effects (42). CNO has a slower onset and longer duration of action than DCZ (43), potentially better mimicking the timescale of a clinical anesthetic in neural ensemble activation. However, in both *null* and Gq^+^ mice, there were no significant effects of CNO on the latency to fall compared to saline (**Fig 2E-F**), indicating that chemogenetic stimulation of anesthesia-activated ensembles does not impact motor coordination on the rotarod. See **Table S2** for detailed statistics.

### Chemogenetic stimulation induces anesthetic-like immobility in a dose-dependent manner

As immobility is used as a defining behavioral outcome of general anesthesia (44), we hypothesized that chemogenetic stimulation of isoflurane anesthesia ensembles would decrease general locomotion in Gq^+^ mice. To test this, mice were assessed for locomotor behavior in a modified open field following alternating trials of saline, 0.01mg/kg DCZ, or 0.1mg/kg DCZ administration. Overall, we observed increased immobility after chemogenetic stimulation with DCZ in Gq+ mice (**Fig 3, Movie S2**). *Null* littermates did not show significant differences in their distance moved or velocity between DCZ and saline trials (**Fig 3A-B**). *Null* controls also did not show significant freezing behavior after chemogenetic stimulation (**Movie S2,** see **Table S2** for detailed statistics).

**Fig 3.**
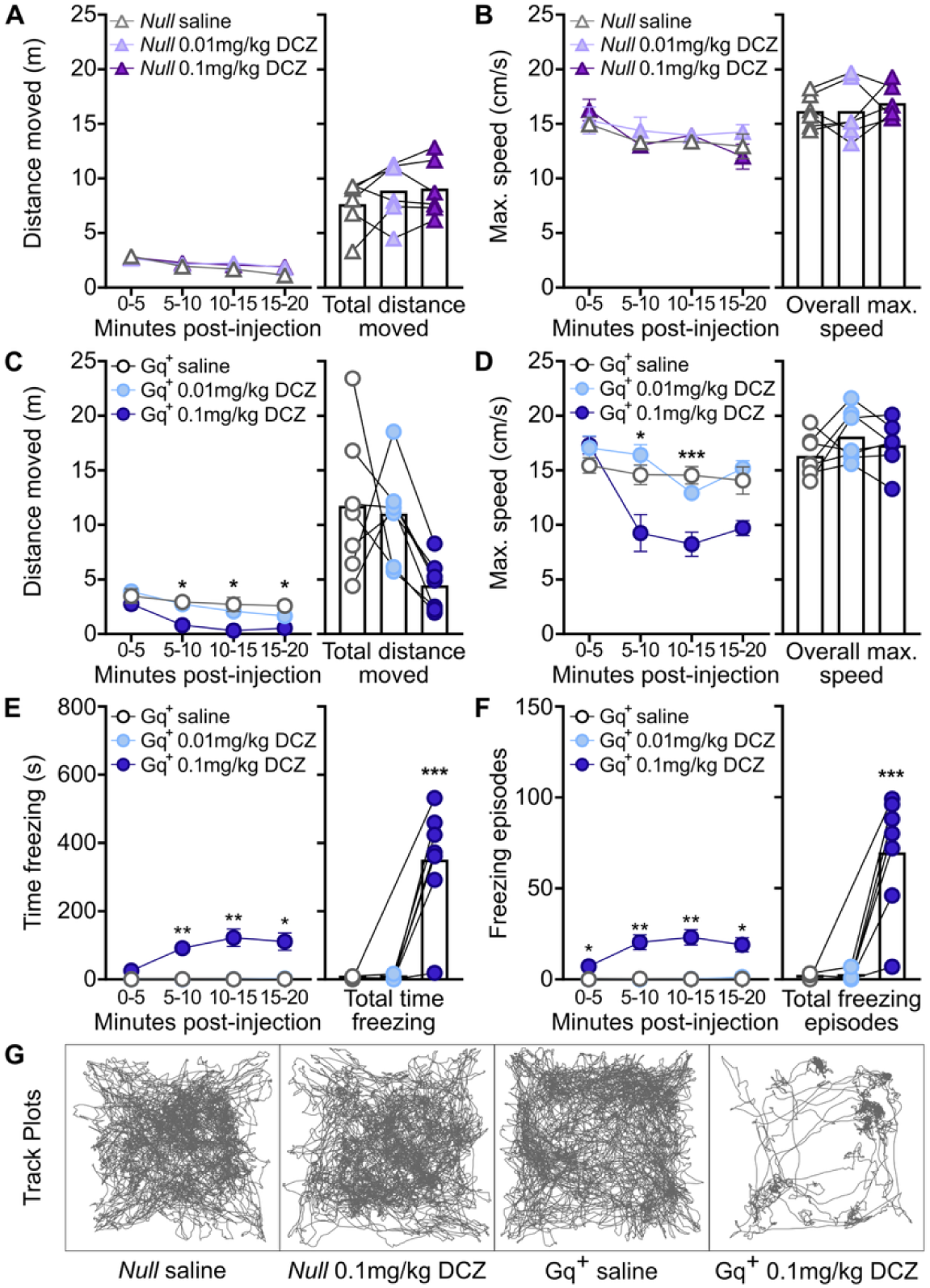
Chemogenetic stimulation with DCZ induces anesthetic-like immobility in a dose dependent manner. Compared to saline, there were no significant effects of either DCZ dose on **A)** distance moved or **B)** maximum speed in *null* littermate controls (n=6) in a modified open field. Gq^+^ mice (n=7) showed a significant decrease in **C)** distance moved over time and **D)** maximum speed over time following injection with 0.1mg/kg DCZ, compared to saline. Gq^+^ mice also showed a significant increase in **E)** time spent freezing and **F)** freezing episodes in a modified open field following injection with 0.1mg/kg DCZ, compared to saline. Representative track plots generated by ANY-maze are shown in **G)**. Error bars represent standard error of the mean.*p<0.05, **p<0.01, ***p<0.001 vs. saline. For detailed statistics see **Table S2**.

Gq+ mice showed dose-dependent decreases in distance moved and velocity after chemogenetic stimulation of anesthesia-activated ensembles (**Fig 3C-D**). Distance moved was significantly lower after 5 minutes post-injection with 0.1mg/kg DCZ compared to saline. Neither DCZ dose significantly affected the total distance moved (**Fig 3C**). Chemogenetic stimulation with 0.1mg/kg DCZ decreased maximum speed between 5-15 minutes post-injection in Gq^+^ mice, with no decrease in overall maximum speed (**Fig 3D**).

Notably, Gq^+^ mice showed a significant, dose-dependent increase in the time spent freezing following injection with DCZ (**Fig 3E**). 0.1mg/kg DCZ significantly increased freezing episodes at every time point and in total (**Fig 3F**). Representative track plots of distance moved in the modified open field are shown in **Fig 3G**.

### Chemogenetic stimulation with CNO induces anesthetic-like immobility

Given that the effects of DCZ on immobility were not completely sustained for the duration of the test, we performed additional tests of anesthetic-like immobility following chemogenetic stimulation with CNO to approximate sustained ensemble engagement. In *null* littermate controls, there was no effect of CNO on distance moved (**Fig 4A**) or maximum speed (**Fig 4B**) across time points and overall. *Null* littermates did not show significant freezing behavior with saline or CNO (see **Movie S3** and **Table S2** for detailed statistics).

**Fig 4.**
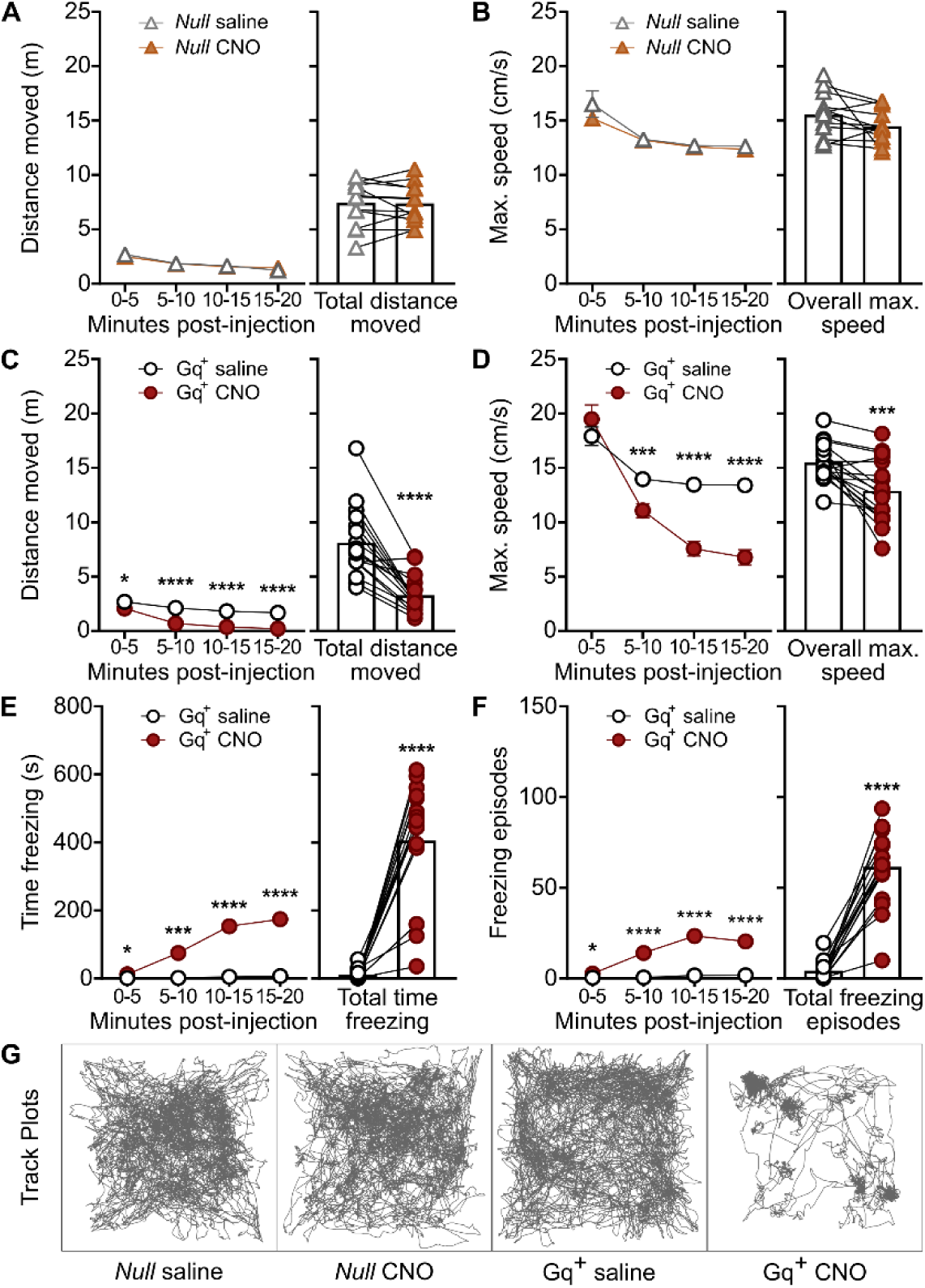
Chemogenetic stimulation with CNO induces anesthetic-like immobility. Compared to saline, there were no significant effects of CNO injection on **A)** distance moved or **B)** maximum speed in *null* littermate controls (n=12) in a modified open field. Gq^+^ mice (n=18) showed a significant decrease in **C)** distance moved and **D)** maximum speed following injection with 5mg/kg CNO, compared to saline. Gq^+^ mice also showed a significant increase in **E)** time spent freezing and **F)** freezing episodes in a modified open field following injection with 5mg/kg CNO, compared to saline. Representative track plots generated by ANY-maze are shown in **G)**. Error bars represent standard error of the mean. *p<0.05, **p<0.01, ***p<0.001, ****p<0.0001 vs. saline. For detailed statistics see **Table S2**.

Gq^+^ mice showed a significant decrease in the distance moved following injection with CNO compared to saline, at every time point and in total (**Fig 4C**). This was accompanied by a significant decrease in maximum speed after 5 minutes post-injection, as well as in overall maximum speed (**Fig 4D**). Gq^+^ mice injected with CNO showed a significant increase in time freezing at all time points, as well as the total time spent freezing, when compared to saline (**Fig 4E**). There was a significant increase in freezing episodes at all time points and overall (**Fig 4F**). All locomotor effects became more pronounced over time resulting in immobility. Representative track plots of distance moved in the modified open field after CNO stimulation are shown in **Fig 4G**.

### Chemogenetic stimulation produces analgesia

Analgesia is another dissociable behavioral outcome of general anesthesia. Some general anesthetics have analgesic effects at subanesthetic doses, or doses that do not produce a loss of consciousness or significant sedation (45). To assess potential effects on analgesia after brain-wide chemogenetic stimulation of isoflurane ensembles, mice were tested on warm water tail withdrawal, an assay of thermal nociception (46). We assessed mice at baseline following saline injection, and then at 30-minute intervals following injection with DCZ (0.01 or 0.1 mg/kg) or CNO (1 or 5 mg/kg, **Fig 5A**, see **Table S2** for detailed statistics). At 30 min following 0.01 mg/kg DCZ, a dose that did not previously produce significant sedation or immobility, Gq^+^ mice showed an increased latency to tail flick from baseline compared to *null* littermates, suggesting anti-nociception. After 0.1mg/kg DCZ, Gq^+^ mice showed an overall greater latency to tail flick (**Fig 5B**). After 1mg/kg CNO, Gq^+^ mice showed a trend for increased latency to tail flick at 30 min (p=0.053) and their latency to tail flick was significantly increased at 60 and 90 min compared to *null* littermates (**Fig 5C**). After 5 mg/kg CNO, Gq+ mice showed a significantly increased latency to tail flick at all time points tested (**Fig 5C**). Together, these data indicate that stimulation of isoflurane-activated ensembles has anti-nociceptive effects that are also observed independent of behavioral sedation.

**Fig 5.**
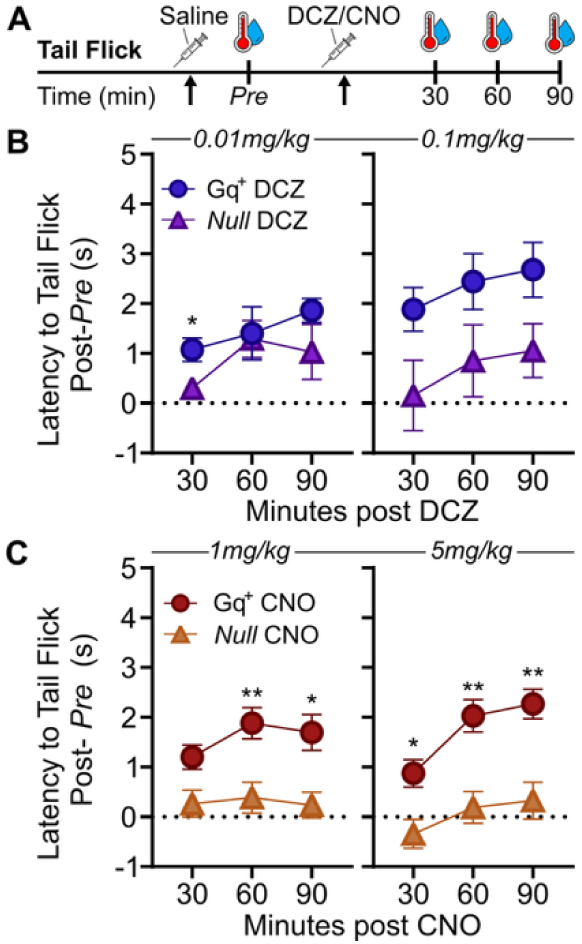
Chemogenetic stimulation produces thermal anti-nociception in Gq^+^ mice. **A)** Mice were tested for their latency to withdraw their tail from a warm water bath (tail flick). Baseline behavior (*Pre*) was recorded after saline injection on each test day. Immediately afterwards, mice received either 0.01 or 0.1mg/kg DCZ, or 1 or 5 mg/kg CNO. Latency to tail flick was tested again at 30, 60, and 90-min post-DCZ or CNO. **B)** Gq^+^ mice (n=11) showed an increase in latency to tail flick compared to *null* littermates (n=6) at 30-min post injection with 0.01mg/kg DCZ, but not at following timepoints. 0.1mg/kg DCZ induced a general increase in latency to tail flick in Gq^+^ mice compared to *null* littermates. **C)** Gq^+^ mice (n=13) showed significantly increased latency to tail flick compared to *null* littermates (n=12) at 60 and 90-min post-1mg/kg CNO. Gq^+^ mice also showed significantly increased latency to tail flick at 30, 60, and 90-min post-5mg/kg CNO compared to *null* littermates. Error bars represent standard error of the mean. *p<0.05, **p<0.01 vs. *null* littermate controls. For detailed statistics see **Table S2**.

## Discussion

Using a transgenic mouse system for activity-dependent chemogenetic capture, we identify a spatially distributed brain-wide ensemble engaged by isoflurane anesthesia and show that chemogenetic reactivation of this ensemble in the awake state reproduces dissociable components of the anesthetic state (increased anesthetic sensitivity, immobility, and thermal antinociception) without fully recapitulating loss of consciousness as indexed by LORR. These findings support a circuit-specific model of general anesthesia in which distinct behavioral outcomes are encoded by overlapping but functionally separable sub-ensembles within a distributed network, rather than by uniform suppression of brain-wide activity or by any single discrete locus.

A central observation is that the behavioral components of the anesthetic state dissociated in a DREADD ligand dose-dependent manner. Antinociception emerged at DCZ and CNO doses that produced no detectable sedation or locomotor suppression, whereas immobility and freezing required higher doses, and LORR was not reproduced at any dose tested. A parsimonious interpretation is that the captured ensemble contains functionally distinct sub-populations with different activation thresholds. An antinociceptive component is recruited at low levels of ensemble activation, a sedative-immobility component recruited at higher levels, and a consciousness-gating component that either was not adequately captured or that requires recruitment patterns the chemogenetic strategy cannot reproduce. This dissociation aligns with the broader view that general anesthesia is not a unitary state but a composite of separable neural processes (5), and it parallels observations that some general anesthetics produce analgesia at sub-anesthetic doses (45). Importantly, the absence of CNO effects in *null* littermates across every assay argues that these dissociations reflect true ensemble engagement rather than off-target pharmacology.

The anatomical signatures of the captured ensemble reinforces and extends prior work on the neural substrates of anesthesia. We observe convergence onto canonical arousal-regulating regions (VLPO, intralaminar thalamus, lateral hypothalamus, parabrachial complex, periaqueductal gray, and median raphe), indicating that activity-dependent capture under isoflurane successfully tags structures with established causal roles in regulating consciousness and anesthetic sensitivity (13, 21, 22, 26, 47–50). Broad engagement of thalamocortical circuitry, including intralaminar and association thalamic nuclei alongside multiple layers of sensory, motor, and association cortices, is consistent with prior reports that isoflurane disrupts thalamocortical and cortico-cortical communication thought to underlie conscious awareness (23–25, 51, 52). At the same time, this brain-wide transgenic capture strategy revealed regions less emphasized in circuit-level studies of unconsciousness. These include the subicular and parahippocampal structures rather than CA1/CA3 proper, primary auditory and visual cortices, and cerebellar cortex. Together, this suggests that the network engaged during isoflurane anesthesia extends beyond classical arousal circuitry. Whether these regions are functionally required for a specific component of the anesthetic state, or whether they reflect network-level recruitment downstream of direct anesthetic targets, remains an open question that cannot be resolved with this transgenic system.

Several considerations may explain why chemogenetic reactivation did not fully reproduce LORR. First, FosTRAP2 captures approximately 60–70% of Fos-positive neurons (53). This incomplete labeling efficiency may be particularly consequential for a behavior like LORR that may depend on coordinated recruitment of a near-complete circuit, rather than partial recruitment. Second, our 3-hour anesthetic exposure with 4-OHT delivered at the 90-min midpoint preferentially captures neurons active during maintenance of the anesthetic state rather than during induction or the transition into unconsciousness. Thus, the ensemble required to generate LORR may be transiently active during induction and underrepresented in our labeling window. Third, accumulating evidence indicates that non-neuronal cells, particularly microglia, facilitate and sustain anesthesia-induced unconsciousness (54–56), and the activity-dependent strategy used here may not engage these cell populations.

Activity mapping cannot distinguish neurons that directly mediate anesthetic effects from those that are indirectly activated by network-level changes, and isoflurane is known to act on a broad array of receptor systems across the brain (57). This strategy identifies neurons engaged during anesthesia but does not establish which populations are functionally necessary for any given behavioral outcome. The overlap between regions implicated in antinociception (basolateral amygdala, lateral hypothalamus, ventral hippocampus (34–36, 58–62)) and those implicated in arousal regulation means that our chemogenetic stimulation cannot dissociate the contribution of individual nodes to specific behavioral components. Lastly, we used both male and female mice, and although sex distributions are reported, the study was not powered to detect sex-specific effects. This is a relevant consideration given documented sex differences in anesthetic sensitivity (40).

In summary, these results establish this mouse transgenic system as a platform for dissecting the brain-wide neural circuit architecture of anesthesia-induced behavioral states. We provide a cellular resolution map that defines a candidate network with discrete individual nodes that can be interrogated using intersectional genetic approaches, projection-specific manipulations, or combined chemogenetic-electrophysiological readouts to determine which sub-circuits are necessary for which components of the anesthetic state. Extending this strategy to other anesthetic agents would test whether distinct anesthetics converge on shared circuit substrates or recruit divergent networks. More broadly, the behavioral dissociations we observe between unconsciousness, immobility, and analgesia supports a framework in which the clinical goals of general anesthesia could be targeted independently through circuit-specific interventions.

## Materials and Methods

### Animals

All experiments were approved by the National Institutes of Health and the University of Washington Institutional Animal Care and Use Committee. Heterozygous R26-LSL-Gq-DREADD^+/-^ mice (purchased from Jackson Laboratories, stock #026220) (18, 19) were crossed with homozygous Fos^2A-iCreER^ (Fos-TRAP2) mice (17) to generate R26-LSL-Gq-DREADD^+^xFos2a-iCreER^+^ (Gq^+^) and R26-LSL-Gq-DREADD^-^xFos2a-iCreER^+^ (*null*) FosTRAP2 mice. Genotyping was performed by Transnetyx using tail clippings obtained at weaning. Male and female F1 progeny were used for experiments (*null* n=6-12; Gq+ n=7-18). Animals were group-housed in the vivarium under a 12-hour reverse light/dark cycle, with *ad libitum* access to food and water.

### Activity-dependent chemogenetic labeling (TRAP) under anesthesia

At approximately 8 weeks old, Gq^+^ and *null* FosTRAP2 mice were induced with isoflurane (2-3%). Mice were maintained on isoflurane anesthesia (1.2-1.3%) for 3 hours. At the 90-minute (min) halfway point, all mice were injected with 4-hydroxytamoxifen (4-OHT, 50mg/kg, Sigma Aldrich) to induce *Cre*-recombination for neural activity-dependent labeling (63). All mice were immobile and had confirmed loss of reflex. Mice were actively warmed on a heating pad and continuously monitored by an experimenter. After TRAP, all mice recovered for 1 week in the home cage to allow for DREADD protein expression in anesthesia-activated ensembles prior to behavioral testing.

### Loss and return of righting reflex

To assess sensitivity to isoflurane following chemogenetic stimulation, all mice were injected with deschloroclozapine (DCZ, 0.1mg/kg), or saline vehicle, in alternating trials. Five min post-injection, mice individually underwent stepwise isoflurane exposure from 0.2-2.2% in 0.5% increments. Each step was maintained for 2 min and the isoflurane concentration at which loss of righting reflex (LORR) occurred was recorded. The order of DCZ or saline injection was counterbalanced in each group.

### Rotarod test

Mice were habituated to the rotarod at 4 rotations per min (RPM), 10-min daily for two days. Following habituation, mice were tested on the rotarod after saline injection, starting at 4 RPM and accelerating at 2 RPM/min to a maximum of 24 RPM over a 10-minute trial. Mice were then injected with DCZ (0.1mg/kg) or clozapine-n-oxide (CNO, 5mg/kg) and tested again 10 min post-injection. Latency to fall was recorded, with a maximum duration of 10 min per trial.

### Modified open-field test

Mice were habituated to a modified small open field chamber (15x15x15cm) for 3 days. Mice were placed individually in the modified open field after injections of saline, DCZ (0.01 and 0.1 mg/kg), or CNO (5 mg/kg) in alternating trials and allowed to explore freely for 20 min with concurrent video recording. There was a minimum 24-hour washout between CNO and subsequent saline trials. Distance moved (m), maximum speed (cm/s), total time freezing (s), and number of freezing episodes were analyzed using ANY-maze 7.4 software. Standard sensitivity (slight movement acceptable) was used to detect freezing, with a minimum bout duration of two seconds per freezing bout.

### Tail withdrawal test

Thermal anti-nociception was assessed by the latency for mice to withdraw their tail after 1/3 – 1/2 of the tail is submerged in a warm water bath (52.5°C) per published studies (46, 64, 65). Testing was performed at baseline immediately following saline injection, and at 30, 60, and 90-min following DCZ (0.01 and 0.1 mg/kg) or CNO injection (1 and 5 mg/kg CNO).

### Tissue collection

Following behavioral testing, mice were deeply anesthetized with sodium pentobarbital (100mg/kg) and underwent transcardial perfusion with phosphate-buffered saline followed by 10% formalin. Whole intact brains were stored at 4°C in 0.01M phosphate-buffered saline (PBS) with 0.01% sodium azide until immunolabeling and clearing.

### Whole brain clearing and immunolabeling

A modified version of the iDisco+ procedure was used to immunolabel and clear intact brains as previously published (66, 67). The following antibodies were used: rabbit anti-HA-Tag antibody (Cell Signaling Technology, 1:2500) followed by donkey anti-rabbit AlexaFluor 647 Fab2-conjugated secondary antibody (Jackson ImmunoResearch, 1:500). Please see Madangopal *et al.*, 2022 (67) for a detailed protocol.

### Intact brain imaging with light sheet microscopy

Intact cleared, immunolabeled brains were immersed in ethyl cinnamate and imaged in the horizontal orientation with a light sheet microscope (SmartSPIM, Life Canvas Technologies) using the 3.6X objective. Images were acquired in two channels (488 nm: autofluorescence; 639 nm: mCherry/AlexaFluor-647) and stitched using the SmartSPIM acquisition software. ArivisPro software (Zeiss) was used to digitally re-slice stitched image files to the coronal orientation. Representative z-stacks (150 images) and 3-dimensional images were visualized using Arivis Pro software (Zeiss).

### Analysis of brain-wide expression

Images were down sampled (0.5x) prior to image processing. We trained a pixel classifier in Ilastik to segment single cells based on somatic signal classification and applied a modified ClearMap automated quantification pipeline as previously published (67, 68). This generated segmented cell counts per brain region and voxelized heatmaps. We computed TRAP+ cell counts within two different levels of hierarchy within the Unified Brain Atlas: (1) 9 major anatomical divisions (**Fig 1C**) and (2) 917 minor subregions within those major anatomical divisions that no longer split into further daughter regions (**Table S1**). We provide the raw TRAP+ cell counts, cell density, and statistical results for all analyses (**Table S1**).

### Statistical analysis

Data were checked for outliers using the Grubbs outlier test, and for normality using Shapiro-Wilk. Paired and unpaired t-tests, one-way ANOVA, two-way repeated measures ANOVA with Sidak’s test for multiple comparisons, or one-way repeated measures ANOVA with Dunnett’s test for multiple comparisons were performed using GraphPad Prism. FDR correction was applied to whole brain quantification. Statistical significance was set at alpha < 0.05, q < 0.05 (major anatomical divisions) or < 0.1 (minor subregions). Data is presented as mean ± standard error of the mean. Detailed statistical tests and results are provided in **Tables S1-2**. Raw data are available upon request.

## Acknowledgments

This project was funded by the National Institute of General Medical Sciences R35GM146751 (to MH), T32GM086270 (to MHCN), and supported by the University of Washington Department of Anesthesiology & Pain Medicine. We thank Dr. Eric Szelenyi for his technical support. We thank Dr. Richard Palmiter for sharing Fos-TRAP2 transgenic mice.

## Supplementary Tables

**Table S1.** Brain-wide cell count and density data for Gq-DREADD-expressing neurons across major anatomical divisions and subregions.

**Table S2.** Statistical analyses for all behavioral experiments.

## Supplementary Figures

**Fig S1.**
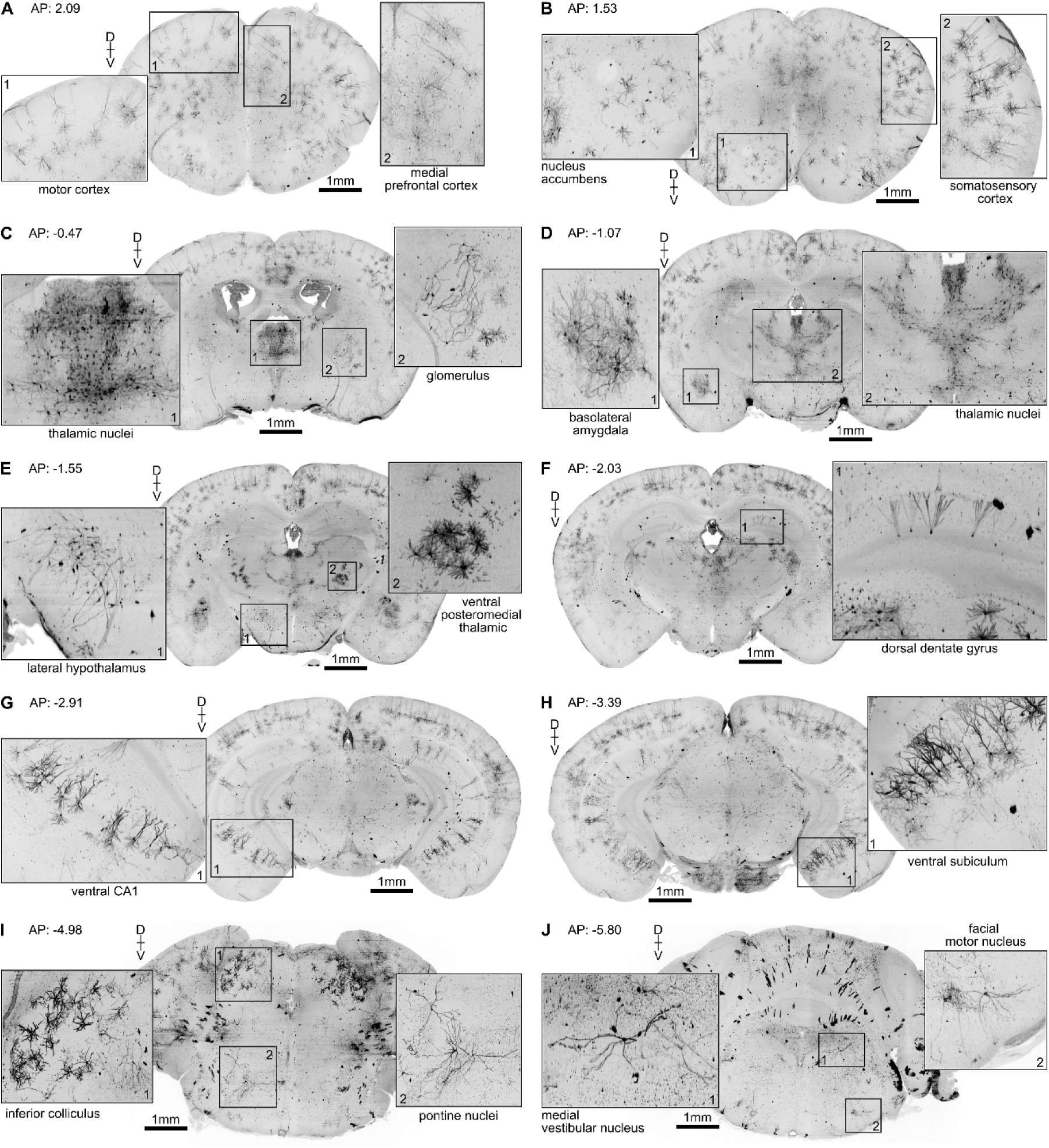
Representative images of DREADD expression across the intact brain in a Gq+ mouse, visually re-sliced in the coronal plane after light sheet microscopy. **A**) 1-motor cortex, 2-medial prefrontal cortex **B**) 1-nucleus accumbens, 2-somatosensory cortex **C)** 1-thalamic nuclei, 2-glomerulus. **D**) 1-basolateral amygdala, 2-thalamic nuclei **E**) 1-lateral hypothalamus, 2-ventral posteromedial thalamic **F)** dorsal dentate gyrus **G)** ventral CA1 **H)** ventral subiculum **I)** 1-inferior colliculus, 2-pontine nuclei **J)** 1-medial vestibular nucleus, 2- facial motor nucleus. Scale bar 1 mm.

## Supplementary Movies

**Movie S1.** DREADD expression across the intact brain visualized with light sheet microscopy.

**Movie S2.** Representative videos of *null* and Gq+ mice in the modified open field following treatment with saline or 0.1mg/kg DCZ.

**Movie S3.** Representative videos of *null* and Gq+ mice in the modified open field following treatment with saline or 5mg/kg CNO.

